# Dopamine ramps as a normative consequence of dual-process control

**DOI:** 10.64898/2026.02.18.706579

**Authors:** Luke Priestley, Thomas Akam

**Affiliations:** Department of Experimental Psychology, University of Oxford, Oxford, UK; Sainsbury Wellcome Centre, University College London, London, UK

## Abstract

Midbrain dopamine neurons are thought to implement a temporal difference (TD) reward prediction error (RPE) that updates cached values stored in striatum. This has been challenged by evidence that dopamine “ramps up” to predictable rewards during goal-directed behaviour. Here, we propose that dopamine ramps are RPEs generated by a dual-process learning system in which values inferred using a world model train cached values via the RPE. Ramps arise because efficient training of cached values requires that inferred values contribute to the update target but not the prediction component of the RPE. The model reproduces key dopamine ramp phenomena, including learning dynamics on fast and slow timescales, global updates following changes in reward expectation, transient responses during unexpected state transitions, and sensitivity to state uncertainty manipulations. We therefore argue that dopamine ramps are a signature of interactions between inferred and cached values that revise the traditional dichotomy between model-based and model-free learning.

## 2 Introduction

Dopamine is widely thought to implement a temporal-difference (TD) reward prediction error (RPE) signal that updates cached values stored at striatal synapses (Montague et al., 1996; Schultz, 2006; Schultz et al., 1997). However, there are features of dopamine activity that this theory struggles to explain. Perhaps most striking is the fact that dopamine in ventral striatum “ramps up” in anticipation of predictable rewards (Howe et al., 2013), particularly in spatial paradigms where distinct locations or sensory states indicate reward proximity (Farrell et al., 2022; Guru et al., 2020; Hamid et al., 2016; Kim et al., 2020; Krausz et al., 2023; Mikhael et al., 2022; Mohebi et al., 2019). The tension with the RPE hypothesis is clear: if dopamine implements an RPE, why do dopamine signals progressively increase during goal approach in well-learned tasks when the value of each state is known?

Theoretical accounts of dopamine ramps fall broadly into two camps. The first proposes that dopamine conveys a value signal rather than an RPE (Hamid et al., 2016; Howe et al., 2013; Mohebi et al., 2019). Although this explains why dopamine might ramp in spatial tasks, where value increases with reward proximity, it is difficult to reconcile with evidence that dopamine exhibits key properties of an RPE in many species and settings (Blanco-Pozo et al., 2024; Eshel et al., 2015; Kim et al., 2020; O’Doherty et al., 2003; Pessiglione et al., 2006; Schultz et al., 1997; Steinberg et al., 2013; Witten et al., 2011). The second proposes that dopamine ramps are a specific case of RPE that arises under special conditions – for example, when state-uncertainty prevents accurate value estimation (Mikhael et al., 2022), when synaptic decay induces forgetting (Kato and Morita, 2016), or when action-timing is uncertain (Lloyd and Dayan, 2015).

Two, striking, recently reported features of dopamine ramp dynamics are not captured by existing theories. First, new information about expected reward at navigational goals rapidly and globally updates ramp amplitude in a manner that appears inconsistent with TD learning (Guru et al., 2020; Krausz et al., 2023). Second, ramps diminish with experience when animals navigate to the same goal in a stable environment (Guru et al., 2020), but on a much slower timescale than that over which behaviour converges. Guru et al. (2020) and Krausz et al. (2023) propose that model-based value computations underlie the rapid effect of new reward information on dopamine ramps, but a theoretical account explaining why model-based computations give rise to dopamine ramps, and how this explains observed ramp dynamics, is lacking.

Here, we suggest that dopamine ramps are a consequence of a dual-process learning architecture that combines model-based and TD learning systems. This builds on the longstanding idea that the brain has multiple complementary learning systems: an efficient but slow TD system for learning cached values, implemented in the basal ganglia, and a flexible but constrained modelbased system for inferring value using a world-model, putatively implemented in frontal cortex (Balleine and Dickinson, 1998; Daw et al., 2005; Dolan and Dayan, 2013). Our key claim is that if the brain can infer values independently of the TD system, for these to efficiently train cached values they must enter the RPE computation in a specific way that necessarily generates ramps. Specifically: inferred values should contribute to the update target towards which cached values are incremented, but not to the prediction against which outcomes are compared to compute the RPE, which should be determined by cached values alone.

We show that incorporating inferred values into the RPE in this way is normative in the sense that it accelerates learning compared to alternative approaches. We further show that the model generates RPE ramps analogous to dopamine ramps observed in experiments, and reproduces diverse experimental findings on dopamine ramp dynamics. We therefore argue that dopamine ramps are RPEs generated by a normative dual-process learning system, a view that revises the traditional dichotomy between model-free and model-based evaluation.

## 3 Results

We first review the standard TD learning algorithm, its putative implementation in the brain, and its inconsistency with dopamine ramps. The TD algorithm aims to learn a value function that reflects expected cumulative future reward given a starting state and policy (Sutton and Barto, 2014). Formally:

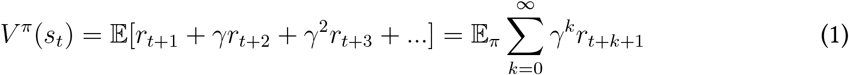

Where *r*_*t*_ is a reward received at time *t, γ* is a discount factor, and *π* is a policy specifying a probabilistic mapping between states of the environment and actions by the agent. For simplicity, we henceforth omit the *π* superscript from all notation.

If the environment is Markovian, *V* (*s*_*t*_) can be expressed recursively as:

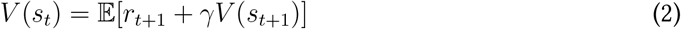

The TD algorithm exploits this recursion using an online learning rule where value estimates are updated in light of immediate rewards, and differences in value between successive states. This is formalised in a teaching signal called the reward prediction error *δ*_*t*_, defined as:

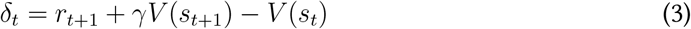

In this equation, the estimate of the old state’s value *V* (*s*_*t*_) is a prediction – i.e., it is the current ‘best guess’ about expected future reward – while the immediate reward *r*_*t*+1_ plus the discounted value estimate for the new state *V* (*s*_*t*+1_) is an update target – i.e. the value toward which the prediction is adjusted. Value estimates are updated using RPEs as:

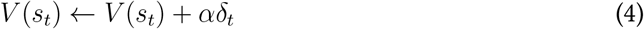

Where *α* ∈ [0, 1] is a learning rate controlling how value estimates change in response to RPEs. The TD algorithm is the basis of an influential account of value-learning in cortico-striatal circuits (Montague et al., 1996; Schultz et al., 1997). This account has three main components fig. 1A-i: (i) The cortex constructs a state representation from sensory experience and communicates it to the striatum (Chang and Tsao, 2017; Liu et al., 2016; Yamins and DiCarlo, 2016); (ii) the striatum estimates value using cortico-striatal synaptic weights, which reflect the relationship between state-features and value (Samejima et al., 2005; Van Der Meer, 2009), and; (iii) VTA dopaminergic neurons compute reward prediction errors that induce plasticity at cortico-striatal synapses (Pawlak and Kerr, 2008), enabling stored values to be updated.

**Figure 1:**
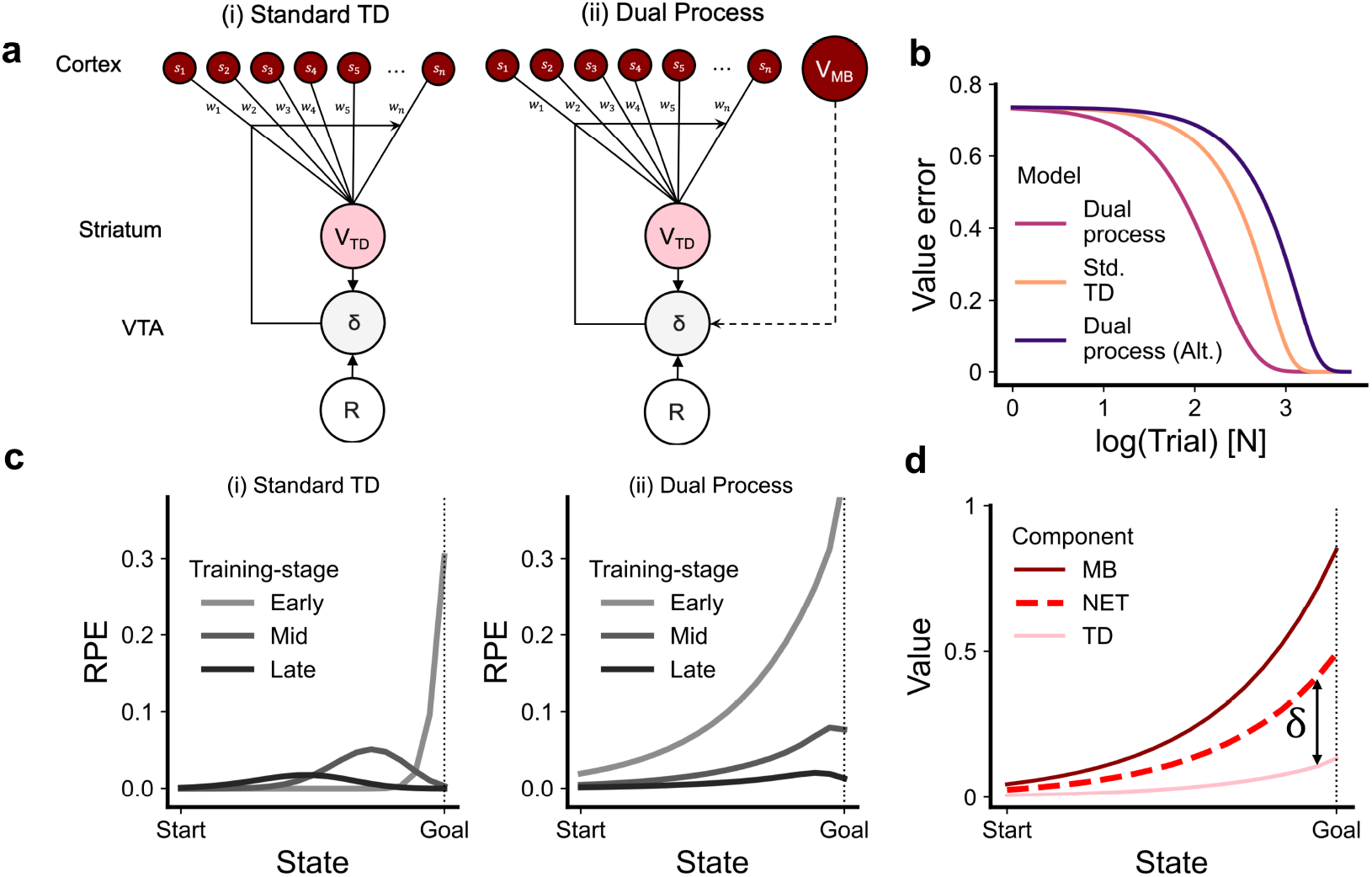
Inferred values train cached values in a dual process architecture. **(a)** Diagram of neural circuits for value-learning showing: (i) The standard model of TD learning in cortico-striatal circuits. (ii) The proposed dual-process model. Note the connection from a model-based evaluation system in frontal cortex to dopamine neurons that bypasses striatum. **(b)** Difference between cached values and the true value function (i.e., value error) during learning in the dual-process model, a standard TD agent, and an alternative dual-process model where *V*_*NET*_ contributes to both the RPE update target and prediction. **(c)** RPEs during approach to a reward at different stages of learning (early, mid, late) in: (i) a standard TD agent, and; (ii) the dual-process model. **(d)** Value functions in different components of the dual-process model during approach to a reward early in learning. The RPE (*δ*) is approximately the difference between *V*_*TD*_ and *V*_*NET*_.

Dopamine ramps challenge this account because the TD algorithm does not, in general, produce RPE ramps in settings where dopamine ramps occur. This is because cached values are assumed to control the agent’s policy. In stable and deterministic environments, cached values converge on the true value function with learning, implying that the policy converges when RPEs disappear. This is inconsistent with experimental observations of dopamine ramps, which persist long after animals exhibit expert task performance (Howe et al., 2013; Krausz et al., 2023).

### 3.1 Inferred values train cached values in a dual process architecture

We propose a dual-process account of dopamine ramps with two core assumptions: (i) that the brain possesses a model-based system that can infer value independently of the basal ganglia TD system in tasks where dopamine ramps occur, and; (ii) that inferred value estimates contribute to training the TD system (fig. 1A-ii). There are many proposals for how the brain might implement model-based value computations (Akam and Walton, 2021; Dolan and Dayan, 2013; Mattar and Lengyel, 2022), including roll-out-based planning, successor or geodesic representations (Dayan, 1993; Sagiv et al., 2025), and inference mechanisms based on attractor dynamics (Donnarumma et al., 2025; Jensen et al., 2025). We do not provide a substantive account of model-based evaluation in this paper. For simulation purposes, we assume a model-based system that infers a goal-conditioned value function using shortest-path distances between states (see below).

If the model-based system can predict future rewards that are not yet reflected in cached values, it can be used to train cached values via the RPE. We argue that to accomplish this, inferred values should contribute to the RPE as:

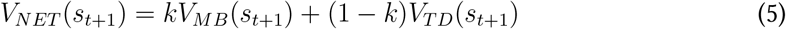

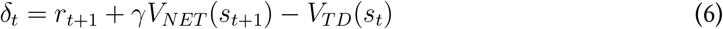

Here, the estimated value of the new state *V*_*NET*_ (*S*_*t*+1_) used in the update target combines an inferred value estimate *V*_*MB*_ and a cached value estimate *V*_*TD*_ using a mixing parameter *k*. Although our model is agnostic as to how *k* is determined, it should in principal ensure that *V*_*NET*_ reflects the agent’s best estimate of the true value given the reliability of *V*_*TD*_ and *V*_*MB*_ e.g., using uncertainty- or confidence-based arbitration (Daw et al., 2005; Lee et al., 2014). We do not provide a substantive account of arbitration in this paper and our simulations use a fixed value of *k* = 0.5 for simplicity (see Methods). Cached values are then updated using the RPE as:

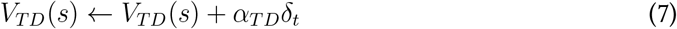

Where *α*_*TD*_ is a learning-rate for cached values.

The critical feature of the RPE computation in eq. (6) is that inferred values contribute only to the update target, as *V*_*MB*_(*s*_*t*+1_), whereas cached values contribute to both the update target as *V*_*TD*_(*s*_*t*+1_), *and* the prediction as *V*_*TD*_(*s*). The rationale is that because the RPE functions to update cached values, the prediction against which outcomes are compared should reflect cached values alone. However, the update target which cached predictions are updated towards should represent the best available estimate of future reward, and hence should incorporate inferred value estimates if they are available. Computing RPEs in this way updates cached values towards *V*_*NET*_, which is desirable when *V*_*MB*_ contains accurate value information that is not yet consolidated into cached values.

How might this dual process architecture be implemented in the brain? A key assumption is that inferred values arise independently of striatal cached values but contribute to RPE computations in the VTA. Since frontal cortex is strongly implicated in model-based evaluation (Akam et al., 2021; Daw et al., 2011; Huang et al., 2020; Jones et al., 2012; Killcross and Coutureau, 2003; Niedringhaus and West, 2022; Stalnaker et al., 2014), we propose that monosynaptic projections from frontal cortex to VTA (Babiczky and Matyas, 2022; Beier et al., 2015; Gao et al., 2022; Wang et al., 2020) communicate the inferred value information used in RPE (fig. 1A-ii, see Discussion).

### 3.2 Dual process learning generates ramping RPEs

We first evaluated the dual-process model on a 1D tabular environment with a single, absorbing, rewarding goal-state. This mimics the trial-based structure of experiments where dopamine ramps occur, which involve moving through a series of locations to a rewarding goal. Environments with a single, terminal reward yield a special case of the value function where value reduces to the reward available in the goal state discounted by its shortest-path distance from the current state:

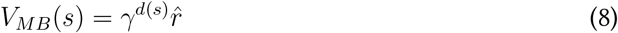

Where *d*(*s*) is the distance between state *s* and the goal-state and 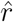 is an estimate of the reward in the goal-state. We assume that distances between states *d* are known *a priori*. In spatial navigation, this assumption is motivated by entorhinal grid-cells which represent 2D space in a manner that generalises between environments and, in principle, permits distance estimation between locations (Bush et al., 2015; Hafting et al., 2005; Whittington et al., 2020). The estimated reward in the goal-state 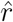 is updated using a delta rule:

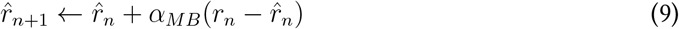

Where *n* indexes a trial of experience, *r*_*n*_ is the reward received on trial *n*, and *α*_*MB*_ is a learningrate for reward estimate updates. We assume that the learning-rate in the model-based system is greater than the learning-rate for the TD system (*α*_*MB*_ *> α*_*TD*_).

We tested the dual process model’s learning performance by comparing time-to-convergence for *V*_*TD*_ with respect to the true value function in three different cases: (i) the dual process model proposed in eqs. 5—7, where *V*_*MB*_ contributes to the RPE update target but not the prediction; (ii) an alternative dual process model where *V*_*MB*_ contributes equally to the RPE update target and prediction, and; (iii) the standard TD learning algorithm, where *V*_*MB*_ does not appear (fig. 1B). Cached values converged to the true value function more efficiently in our proposed model compared to both alternatives. Strikingly, the alternative dual-process model, where *V*_*MB*_ contributed to both the RPE update target and prediction, learned less efficiently than standard TD. This demonstrates that if inferred values are available, it is normative to use them only in the RPE update target.

We next compared RPEs from the dual-process model and standard TD learning during navigation of the linear track environment (fig. 1C). Ramping RPEs occurred in the dual-process model when inferred values predicted future reward that was not yet captured in cached values. We illustrate this in fig. 1D by displaying *V*_*TD*_, *V*_*MB*_, and *V*_*NET*_ during early learning. Inferred values emerged rapidly during initial encounters with the environment, whereas cached values emerged incrementally. Consequently, inferred values exceeded cached values in all states during early learning. Given that *V*_*MB*_ contributes only to the update target, and the prediction is given only by *V*_*TD*_, the RPE is shaped by differences between *V*_*MB*_ and *V*_*TD*_ for successive states, which increase as the agent approaches reward – in other words, RPEs ramp. Dopamine ramps are therefore consistent with inferred values training cached values via the RPE.

### 3.3 Dopamine ramp dynamics at short and long timescales

A key prediction of the dual process model is that dopamine ramps will diminish over time in stable environments. This is because ramps arise from the difference between cached *V*_*TD*_ and inferred *V*_*MB*_ values which, in stable environments, (fig. 1) reduces with experience as *V*_*TD*_ and *V*_*MB*_ converge to the true value function. (fig. 2C). Consequently, RPEs – and therefore ramps – should also reduce with experience. Consistent with this, Guru et al. (2020) report that dopamine ramps in mice gradually diminish with extensive training in a spatial task involving navigation between rewards at alternate ends of a linear track (fig. 2A–B). Dopamine ramps then re-emerged when place-reward contingencies changed, suggesting that ramps implement a learning-related computation consistent with an RPE. RPE ramps in the dual process model reproduced these patterns when simulated on an analogous task (fig. 2D). The dual-process model is thus consistent with long-timescale changes in dopamine ramps reported by Guru et al. (2020).

**Figure 2:**
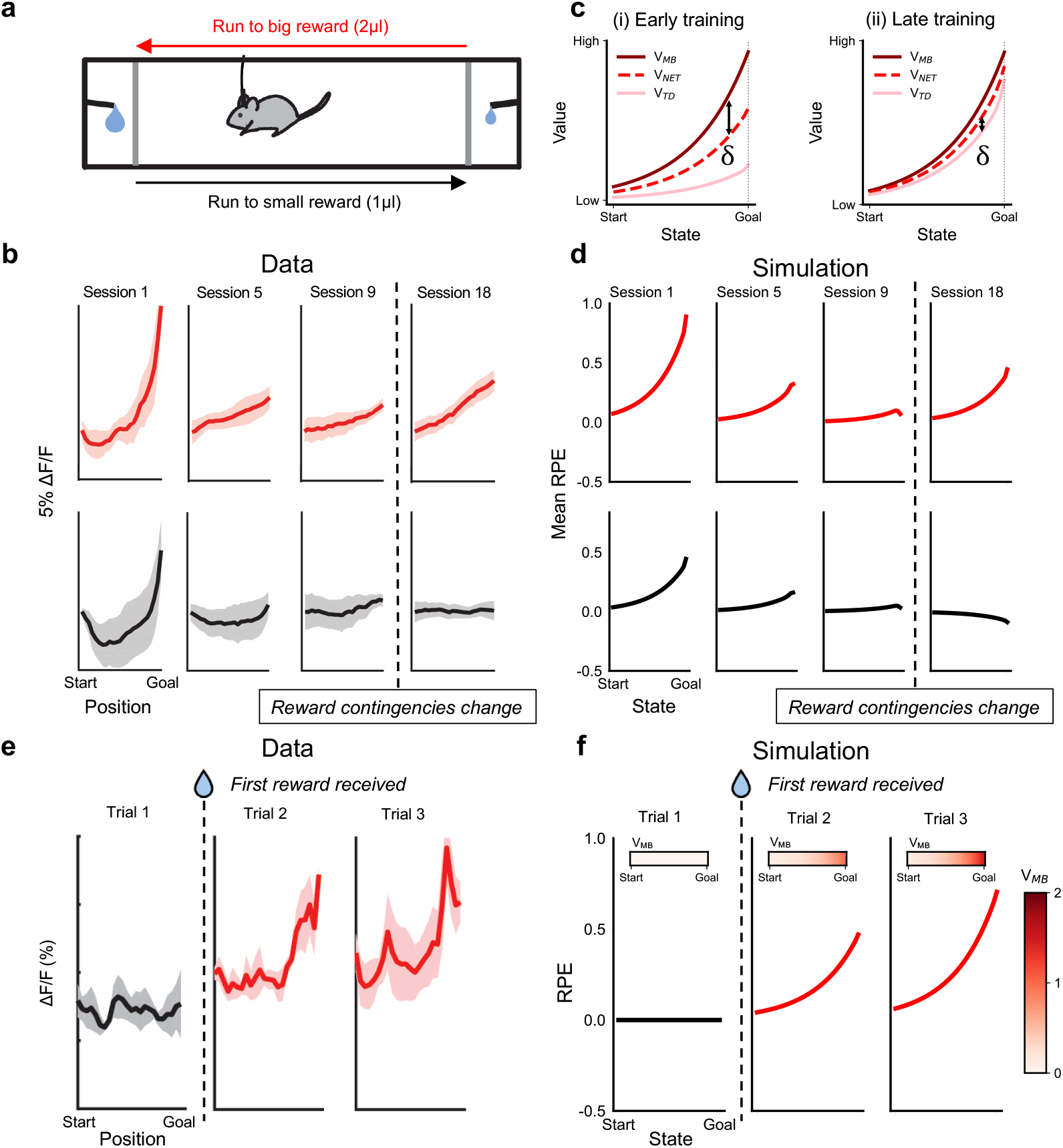
Dopamine ramp dynamics at different timescales. **(a)** Diagram of behavioural task in Guru et al. (2020). **(b)** Experimental data from Guru et al. (2020) showing dopamine ramps at different training stages during runs to big (red) and small (black) reward locations. **(c)** Evolution of value estimates during training in the dual-process model; **(i)** value estimates during early training; **(ii)** value estimates during late training **(d)** Evolution of dual process model RPEs during training on a task that replicates Guru et al. (2020). **(e)** Experimental data from Guru et al. (2020) showing rapid development of dopamine ramps after initial encounters with rewarding goals in a novel environment. **(f)** Evolution of RPEs in the dual-process model during initial encounters with rewarding goals.

Guru et al. (2020) further report that dopamine ramps emerge rapidly when rewards are first encountered in a novel environment, (fig. 2E). This is consistent with our model under the assumption that distances between spatial states can be estimated with minimal experience (see above and discussion), and that the model-based system (*α*_*MB*_) employs a high learning-rate for the reward function. Simulating the dual process model in a novel environment confirmed that RPE ramps were absent on the first trial when the the reward at the goal was unknown. Ramps then rapidly developed over subsequent trials as rewards drive learning of inferred values (fig. 2F). The rapid development of dopamine ramps in novel environments is thus consistent with RPE dynamics in the dual-process model.

### 3.4 Rapid global updates to ramp amplitude by reward

When rewards at goal locations are dynamic, dopamine ramp amplitudes are rapidly updated by reward outcomes. Specifically, Krausz et al. (2023) demonstrate that an outcome at a goal location modulates ramp amplitude on the subsequent visit, with rewards increasing amplitude and omissions decreasing amplitude (fig. 3A–B). Importantly, amplitude changes occur even if the goal is reached by a different route on the subsequent visit, suggesting that they reflect global updates in reward expectation. The dual process model captures these patterns under the assumption that changes in the expected reward at a goal location globally modulate inferred values. This is consistent with inferred values that are computed by combining an estimate of the immediate reward at a goal state with a representation of the distances between states, e.g., via Euclidean distances computed by grid cells (Bush et al., 2015) or cached shortest-path distances as in the Geodesic representation (Sagiv et al., 2025). Implementing the dual process model in a 2D gridworld environment with multiple paths to goal locations recapitulated the patterns reported by Krausz et al. (2023) (fig. 3C–D), suggesting that it is consistent with rapid, global changes in dopamine ramps in environments with dynamic rewards.

**Figure 3:**
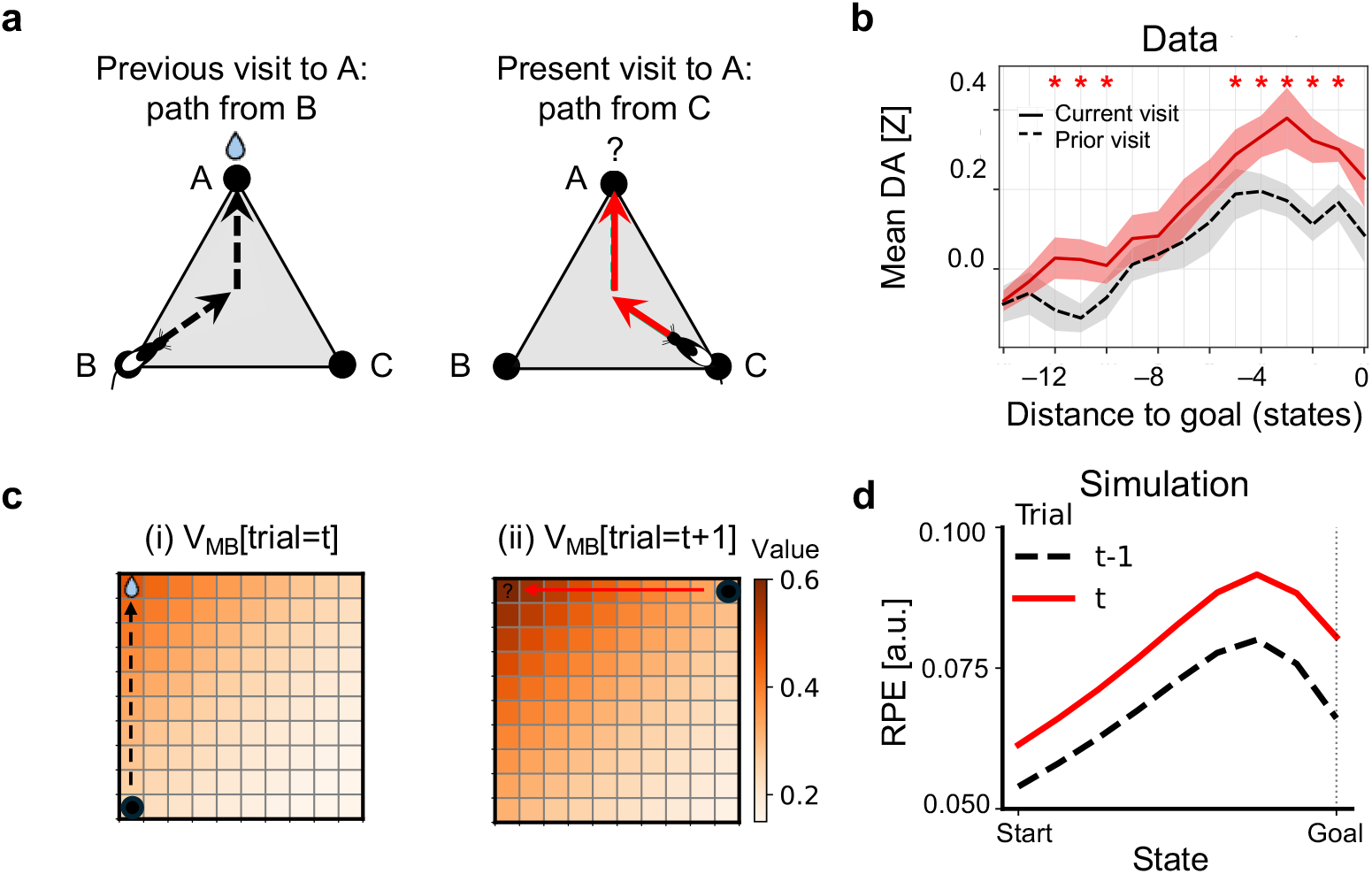
Reward globally updates dopamine ramps. **(a)** Experimental design from Krausz et al. (2023) in which animals can reach rewarding goal locations via multiple routes. **(b)** Experimental data from Krausz et al. (2023) showing that outcomes at goal locations globally update dopamine ramps regardless of subsequent route. **(c)** Diagram showing global updating of inferred values *V*_*MB*_ by rewards in the dual process model. **(d)** Effect of reward on trial *t* on RPEs on trial *t+1* in the dual process model when goal is reached via different routes on *t* and *t+1*.

### 3.5 RPE-like dopamine responses to unexpected state transitions

We next tested whether the dual process model reproduces RPE-like dopamine responses during experimental manipulations in spatial tasks. Previous work has shown that when animals progress toward a reward location in a spatial virtual-reality (VR) environment, dopamine signals are modulated by unexpected changes in spatial position (Kim et al., 2020). For example, teleports between non-adjacent states cause dopamine transients that superimpose on dopamine ramps, with magnitudes that are proportional to teleport end-state (fig. 4A) and teleport-distance (fig. 4B). Similarly, the speed at which animals progress towards the goal modulates ramp slopes, with faster speeds producing stronger slopes (fig. 4C). These patterns favour an RPE interpretation of ramps in which dopamine represents changes in value between timepoints, rather than value itself. Dual process model RPEs reproduced dopamine responses during teleport and speed manipulations because RPEs are intrinsically modulated by changes in value (fig. 4A-C). By equating dopamine with an RPE, the dual process model is thus consistent with the results in Kim et al. (2020).

**Figure 4:**
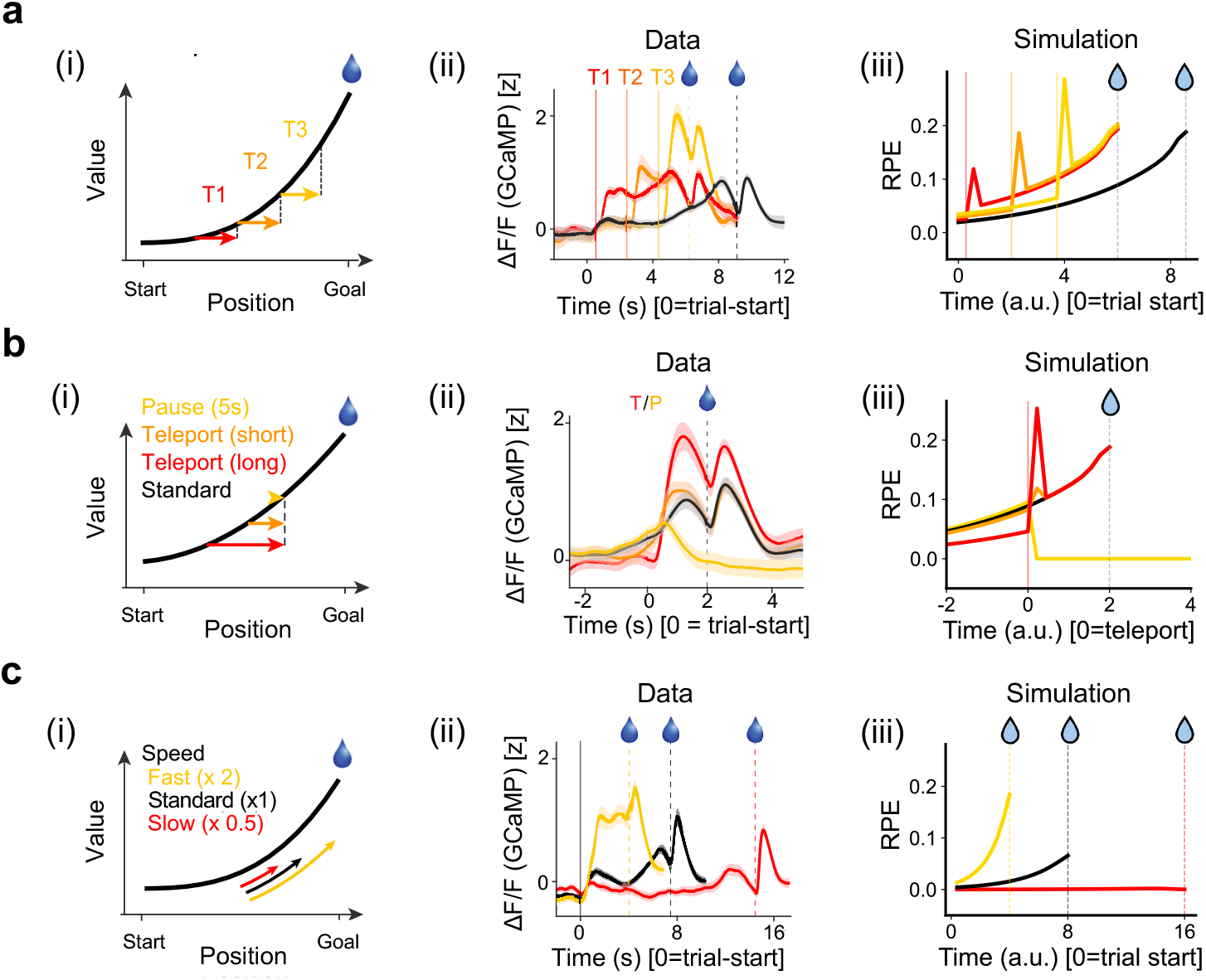
Dopamine responses to unexpected state transitions. Experimental conditions (left), dopamine recordings (middle), and RPEs in simulations of the dual-process model (right) for key experimental conditions from Kim et al. (2020) in which unexpected state transitions occured during reward approach in a VR environment. **(a)** Teleport end-state manipulation, where teleports of constant distance were aligned with different end-states. **(b)** Teleport distance manipulation, where teleports varied in distance but ended at a common state. **(c)** Traversal speed manipulation.

### 3.6 Dopamine ramps dynamics under state-uncertainty manipulations

Finally, we consider experiments motivated by the proposal that dopamine ramps arise from distortions in learning caused by state uncertainty (Mikhael et al., 2022). As in Kim et al. (2020), mice approached a reward location within a VR corridor (fig. 5A). On some trials, the VR environment progressively darkened during goal approach, thereby increasing the animal’s uncertainty about its location (fig. 5A). Dopamine signals on these trials resembled ‘bumps’ rather than ramps – they initially increased more rapidly than the standard trial signal, before subsequently decreasing below it (fig. 5B).

**Figure 5:**
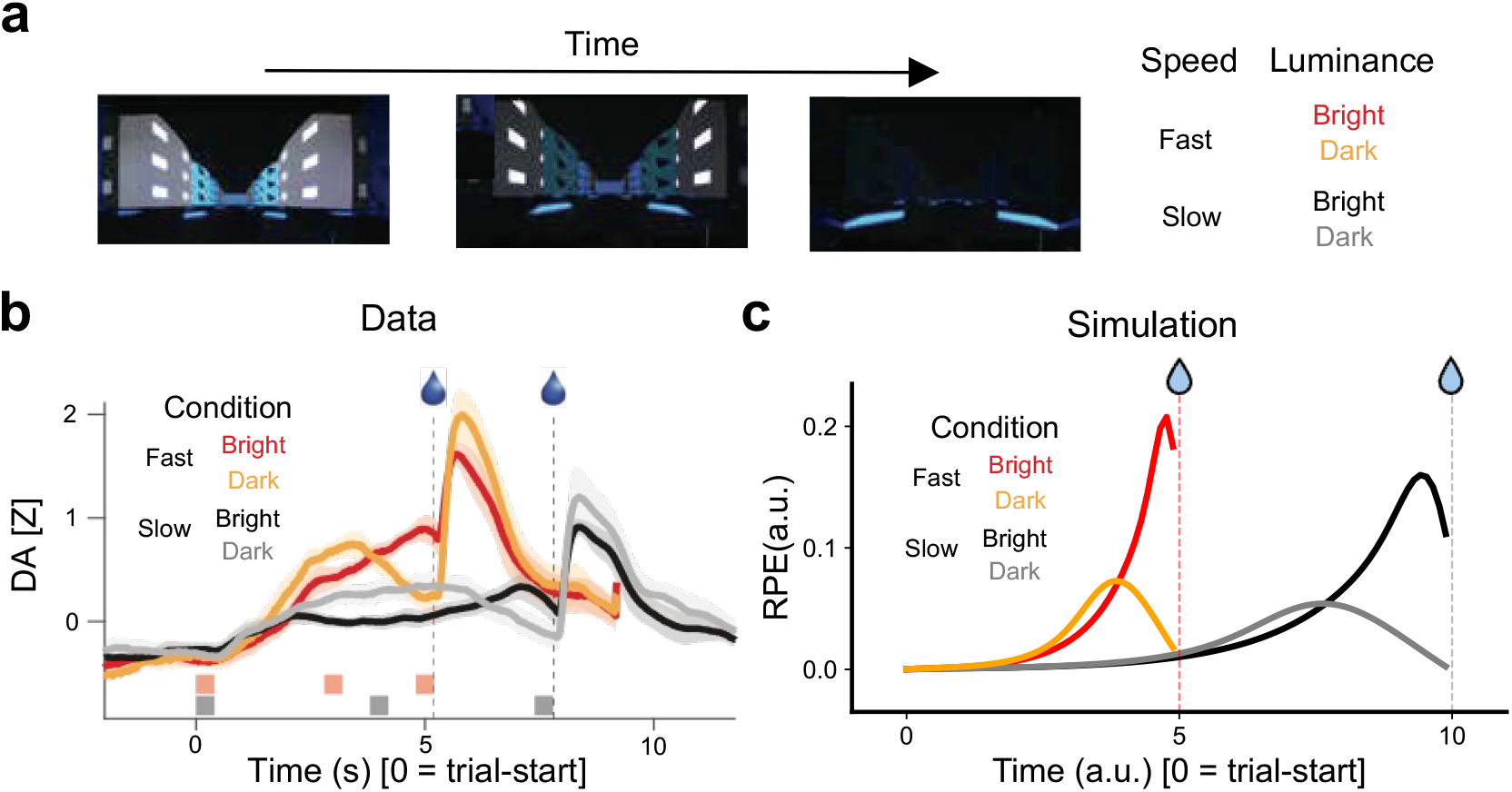
Dopamine ramp dynamics under state-uncertainty manipulations. **(a)** Experimental paradigm from Mikhael et al. (2022) where animals approached a rewarded location in a VR environment that differed in movement speed (fast-vs-slow) and luminance (bright-vs-darkening) across trials. **(b)** Dopamine signals from Mikhael et al. (2022), where progressively increasing state uncertainty causes dopamine bumps rather than dopamine ramps. **(c)** Effect of progressively increasing state uncertainty on RPEs in the dual process model.

Following Mikhael et al. (2022), we assume the subject does not know the true current state *s*_*t*_ of the environment, but instead maintains a probability distribution *p*(*s*| *x*_*t*_) over possible states *s* given sensory input *x*. This probability distribution is assumed to be a Gaussian centred on *s*_*t*_ with standard deviation *σ*_*t*_:

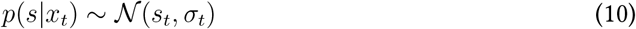

Value estimates with respect to *x* are computed as a probability weighted sum over state value estimates:

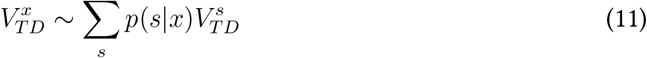

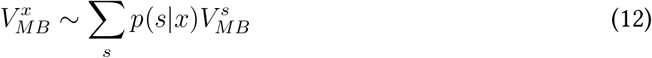

Simulating this version of the model on standard trials with constant state uncertainty produced ramping RPEs consistent with those observed without state uncertainty (fig. 5C). Following Mikhael et al. (2022), darkening trials were modelled by gradually increasing the width of the state uncertainty kernel across the trial. This generated RPE bumps similar to the dopamine bumps seen in the experimental data (fig. 5C).

RPE bumps occur because state uncertainty distorts the inferred value estimates 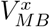 that mdrive RPEs. Greater uncertainty assigns more weight to states away from the true state. Early in the trial, this increases inferred value estimates, as although the uncertainty kernel is symmetric around the true state, the slope of the value function is steeper on the higher value side. Late in the trial, uncertainty adds probability mass primarily behind the true state, as the uncertainty kernel cannot extend beyond the reward location if the reward has not been reached. This decreases inferred value estimates and hence the RPE.

Although state uncertainty manipulations have similar effects in both our model and Mikhael et al. (2022), the underlying mechanism of RPE ramps is fundamentally different. Mikhael et al. (2022) propose that RPE ramps arise from a correction term in the value update which counteracts biases that arise from state uncertainty. Learning therefore converges not when the RPE is zero but rather when it is cancelled out by the correction term. Since the required correction is proportional to state value, this generates RPE ramps. This account rests on the critical assumption that state uncertainty is systematically larger when estimating the value of the new state *V*_*xt*+1_ compared to the value of the old state *V*_*x t*_ in the RPE computation – the rationale being that sensory feedback reduces uncertainty for the old state *s*_*t*_ relative to the new state *s*_*t*+1_. It is this assumption that causes state uncertainty to systematically bias learning, and necessitates the correction term that generates RPE ramps. Importantly, recent experimental data measuring the effect of striatal stimulation on dopamine signals (Campbell et al., 2025) suggests that temporal difference value comparison is implemented by synaptic delays in striatum-VTA circuitry, such that *V*_*x t*_ is simply a delayed copy of *V*_*x t*+1_, and hence inherits the same state uncertainty.

In contrast to Mikhael et al. (2022), our model reproduces dopamine dynamics under uncertainty manipulations without requiring systematic differences in state uncertainty between terms in the RPE computation.

## 4 Discussion

Dopamine ramps have attracted widespread interest because they appear to contradict the theory that dopamine implements a temporal difference reward prediction error (Berke, 2018; Hamid et al., 2016; Howe et al., 2013; Niv, 2013). Here, we propose that dopamine ramps are RPEs generated by a dual-process learning architecture in which values inferred from a world model train cached values via the update target of the RPE. We show that this architecture accelerates cached value learning, and reproduces the key empirical features of dopamine ramps within a unified explanatory framework.

The dual process model generates ramping RPEs because inferred values contribute to the update target but not to the prediction component of the RPE. The rationale is that the update target should represent the best estimate of future reward, and inferred values should therefore contribute to it if they are accurate. The prediction, by contrast, should be determined by cached values alone, because cached values are the quantities that the RPE must update. This asymmetric use of inferred values is normative in the sense that, given accurate inferred values, it accelerates convergence of cached values to the true value function. Strikingly, incorporating inferred values symmetrically in both the update target and prediction slows learning relative to not using inferred values at all (fig. 1C).

The model accounts for key experimental findings on dopamine ramp dynamics via the interplay of two sources of value information (figs. 2 to 4): (i) fast-evolving inferred values, which explain rapid global updates following individual rewards, and (ii) slow-evolving cached values, which explain why ramps diminish with experience as cached values incrementally converge. This clarifies why ramps persist after expert behaviour has developed, as policy can be guided by inferred values long before cached values converge.

Our theory implies that the brain has mechanisms for inferring value online during behaviour, independently of the striatum. This contrasts with offline replay mechanisms that refine cached value estimates guiding subsequent behaviour (Mattar and Daw, 2018; Sutton, 1991). Since modelbased evaluation is generally computationally intensive (Sutton and Barto, 2014), this raises the question of how online inferred value estimation is tractable. We suggest that fast, online inferred value estimation is possible only in specific situations that permit efficient solution methods. Specifically, RL problems characterised by absorbing goal-states reduce goal-conditioned value functions to the immediate reward at the goal, discounted by the distance or cost to reach it. This enables value to be estimated using learned distances between locations (Piray and Daw, 2021; Sagiv et al., 2025). Although real-world behaviour continues after goals are reached, the brain might use the solution methods afforded by absorbing goal states as a heuristic, or as a component of a hierarchical control architecture (Ringstrom et al., 2025).

These considerations suggest that dopamine ramps will emerge when: i) the brain has an internal model of distances between states, and ii) behaviour is organised by discrete, known, and rewarding goal states, rather than random foraging. Goal-directed spatial navigation exemplifies these conditions, explaining the prominence of ramps in navigation tasks. Grid cells facilitate distance estimation in physical space and carry representations that generalize across environments (Bush et al., 2015; Hafting et al., 2005; Whittington et al., 2020), consistent with the rapid onset of dopamine ramps in spatial tasks (Guru et al., 2020). Hippocampal-entorhinal circuits for spatial cognition also represent position in sensory or abstract state spaces (Aronov et al., 2017; Constantinescu et al., 2016), which may explain ramps in tasks where sensory cues indicate reward proximity (Kim et al., 2020). By contrast, classical conditioning tasks lack distinct sensory states indicating reward proximity. Dopamine responses in these settings resemble a backpropagating TD error rather than a ramp, consistent with inferred values playing no role in the update target (Cohen et al., 2012; Schultz et al., 1997).

Neural implementation of the dual process architecture requires that dopamine neurons receive inferred value information through a non-striatal pathway. Although the neural basis of model-based evaluation remains poorly understood, it has been linked to orbitofrontal (OFC) and medial frontal cortex (mFC) in both humans and non-human animals (Akam et al., 2021; Daw et al., 2011; Huang et al., 2020; Jones et al., 2012; Killcross and Coutureau, 2003; Niedringhaus and West, 2022; Stalnaker et al., 2014). Notably, mFC and OFC have monosynaptic projections to VTA dopamine neurons (Babiczky and Matyas, 2022; Beier et al., 2015; Gao et al., 2022; Wang et al., 2020), and stimulating the FC-VTA pathway induces conditioned place preference via the nucleus accumbens (Beier et al., 2015). VTA-projecting FC neurons are therefore a plausible source of the inferred value information that generates dopamine ramps (Guru et al., 2020). Our model further implies that VTA-projecting and striatum-projecting subpopulations of FC neurons should encode different signals: inferred values in the former, and state features in the latter. Consistent with this, these subpopulations are largely anatomically separate, although their coding properties remain uncharacterised (Babiczky and Matyas, 2022; Gao et al., 2022).

Our model makes several testable experimental predictions. (1) VTA-projecting frontal cortex neurons will encode inferred value signals (i.e., expected discounted future reward) in settings where dopamine ramps occur. (2) Silencing the FC-to-VTA pathway, or the components of the world model necessary for inferred value estimation, will abolish dopamine ramps. (3) Abolishing dopamine ramps will slow the development of striatal state-value signals during goal-directed navigation (Van Der Meer, 2009), and alter the development of state value representations during learning. (4) Transiently stimulating FC-to-VTA and striatum-to-VTA pathways will evoke distinct patterns of ventral striatal dopamine release. Stimulating NAc D1 neurons initially excites then subsequently inhibits VTA neurons (Campbell et al., 2025), consistent with the dual role of cached values in the RPE update target and prediction. Stimulating the FC-VTA pathway should, by contrast, evoke VTA excitation alone, since inferred values contribute only to the update target.

Finally, together with Mattar and Daw (2018), our work suggests that the traditional dichotomy between model-based and model-free evaluation should be revised. Each account emphasises that model-based mechanisms have a profound influence on the striatal cached value system – online through dopamine ramps, and offline through replay. This implies that the cached value system should not be viewed as model-free, but rather as a long-term memory system for value that is shaped by both temporal-difference learning and model-based evaluation.

## 5 Acknowledgements

We are grateful to Kris Jensen, Eleanor Spens and Marta Blanco-Pozo for helpful feedback on the manuscript. The work was supported by Wellcome Trust Career Development Award 225926/Z/22/Z. For the purpose of open access, the author has applied a CC BY public copyright licence to any Author Accepted Manuscript version arising from this submission.

## 6 Methods

### 6.1 General simulation details

All simulations were implemented in Python v3.14. For simplicity, the policy for all agents in all simulations was deterministic and involved moving directly to the rewarding goal location. Dual-process agents were simulated according to eqs. 5—7. Task specific environments and parameter choices are described below.

Code for replicating the simulations and generating the manuscript figures is available at: https://github.com/lpriestley/da-ramps

### 6.2 Comparison of value-learning algorithms

To characterise whether the dual-process model accelerated value-learning (fig. 1B), we implemented (i) a dual-process agent, (ii) a standard TD agent, and (iii) an alternative dual-process agent on a linear track environment. The standard TD agents was implemented according to eqs. 3—4. The alternative dual process agent was implemented according to:

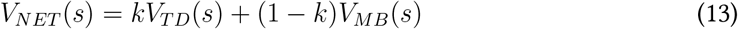

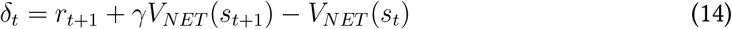

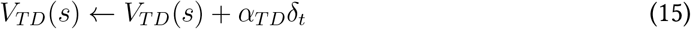

The key difference compared to the dual process agent defined in eqs. 6—7, therefore, is that inferred values appear in both the RPE update target and the prediction, instead of the update target alone. This alternative dual-process agent learned slower than standard TD learning (fig. 1B) and did not generate ramping RPEs. Parameters for the agents were: *α*_*TD*_ = 0.01, *α*_*MB*_ = 0.50, *γ* = 0.93, *k* = 0.50,

The linear track was formalised as a tabular environment with *N* = 10 states. There was a goal state at one end of the track with a scalar reward *r* = 1.0. Agents started each trial at the end of the track opposite the goal state. All agents performed the task for *T* = 5000 trials. Value error was calculated on each trial as 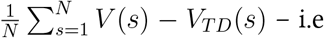. the average discrepancy between cached values *V*_*TD*_ and the true value function *V*.

### 6.3 Characterising dual-process learning dynamics

To demonstrate why the dual-process model produces ramping RPEs, a dual-process agent was simulated on a linear track, formalised as a 1D tabular environment with *N* = 20 states, and a goal state at one end of the track with a scalar reward *r* = 1.0. Agents started each trial at the end of the track opposite the goal state. The parameters for the agent were: *α*_*TD*_ = 0.01, *α*_*MB*_ = 0.50, *γ* = 0.85, *k* = 0.50. The value-functions *V*_*MB*_, *V*_*TD*_ and *V*_*NET*_ were extracted from the agent on trial *t* = 100 of learning and displayed in fig. 1D. RPEs *δ* at early, intermediate and late stages of learning were further extracted and graphed in fig. 1E-ii. This was compared to RPEs from a standard TD agent simulated with parameters *α*_*TD*_ = 0.01, *γ* = 0.85.

### 6.4 Guru et al. (2020) simulation

We compared dual-process RPEs to dopamine signals in Guru et al. (2020). The Guru et al. (2020) study involved recording dopamine signals whilst mice ran between alternate ends of a linear track. One end of the track had a large reward (2*µ*L), and the other end of the track had a small reward (1*µ*L).

To simulate the dual-process model on this task, we treated navigation towards each end of the track as a separate state space, consistent with hippocampal units having strong movement direction tuning on linear tracks. Each agent was simulated with the following parameters: *α*_*TD*_ = 0.005, *α*_*MB*_ = 0.50, *γ* = 0.93, *k* = 0.50. Each linear track was formalised as a 1D tabular environment with *N* = 35 states and a goal state at one end. Agents started each trial at the end of the track opposite the goal state. In the large-reward track, the initial reward value was *r* = 2.0, and the small-reward track, the initial reward value was *r* = 1.0.

Agents performed *S* = 18 sessions of learning, where each session involved *T* = 100 trials. On session *s* = 17, the reward values at the end of high-reward and low-reward tracks were swapped. The training regime was designed to replicate the Guru et al. (2020) experiment.

In fig. 2C, *V*_*MB*_, *V*_*TD*_ and *V*_*NET*_ were extracted from trial *t* = 100 on sessions *s ∈*{1, 4}. In fig. 2D, the evolution of RPEs over learning was visualised by computing, for each state, the mean RPE over trials within a session. In fig. 2F, RPEs and *V*_*MB*_ were extracted for trials *t ∈*{1, 2, 3}.

### 6.5 mKrausz et al. (2023) simulation

We compared dual-process RPEs to dopamine signals in Krausz et al. (2023). In Krausz et al. (2023), rats performed a maze navigation task, in which a series of goal locations delivered probabilistic rewards. The task took place in a complex maze environment with multiple pathways to each goal location.

To simulate the task, a dual-process agent was implemented on a a 2D 10 × 10 gridworld. There was a goal-location *g* = (1, 1) which, when visited, delivered reward stochastically with *r* = 1.0 and *p*(*reward*) = 0.5. The agent was alternately started on odd and even trials from *s*_*odd*_ = (10, 1) and *s*_*even*_ = (1, 10) and followed a trajectory directly to the reward location. This allowed us to test how specific rewards and omissions at the goal location influenced RPEs on the subsequent trial, even when the trajectory to the goal location was different. The agent was simulated with the following parameters: *α*_*TD*_ = 0.01, *α*_*MB*_ = 0.10, *γ* = 0.85, *k* = 0.50. It performed *T* = 500 trials. In fig. 3C, the effect of rewards on inferred values was visualised by extracting *V*_*MB*_ for consecutive trials *t* − 1 and *t*, where *t* − 1 was rewarded. In fig. 3D, the effect of rewards on RPEs was visualised by comparing RPEs on consecutive trials *t* − 1 and *t*, where *t* − 1 was rewarded.

### 6.6 Kim et al. (2020) simulation

We compared dual-process RPEs with dopamine signals in Kim et al. (2020). In the Kim et al. (2020) experiment, subjects viewed a VR track environment with a terminal reward at the end of the track. The environment was manipulated using teleports between non-adjacent states, and speed modulations that controlled how quickly subjects moved through the environment.

We simulated the dual-process agent on these experiments using linear tracks, which were formalised as 1D tabular environments with a goal-state that delivered a reward *r* = 1.0 at one end of the track. The agent always started at the end opposite the goal. In the teleport experiments (fig. 4A and fig. 4B), the track had *N* = 32 states. In the speed-manipulation experiment (fig. 4C), the track had *N* = 40 states. In the teleport-distance experiment, all teleports ended at state *s*_*teleport*−*destination*_ = 24, where short teleports had distance *d*_*short*_ = 2 and long teleports had distance *d*_*long*_ = 10. In the pause condition, the agent remained in *s*_*teleport*_ −_*destination*_ for an arbitrary number of timepoints. We abolished the effect of inferred values on RPEs during the pause period under the assumption that inferred values predict temporally discounted future reward. We assume that such predictions are null in situations when the agent is static in a non-rewarding state. In the teleport end-state experiment, teleports had a constant distance *d* = 10 and were initiated from either early, moderate, or late start locations where *s*_*early*_ = 2, *s*_*intermediate*_ = 10, *s*_*late*_ = 14. In the speed-manipulation experiment, slow, normal and fast speeds were implemented by modulating the step-size with which the agent moved through the environment, where *stepsize*_*small*_ = 1, *stepsize*_*normal*_ = 2, *stepsize*_*fast*_ = 4. Training in the speed-manipulation experiment was performed with *stepsize*_*normal*_. In teleport experiments, the agent was trained on *T* = 200 trials before experiencing the teleport manipulation. In the speed experiment, the agent was trained on *T* = 500 trials before experiencing the speed manipulation. Cached values were clamped during test trials to prevent learning. Agents were simulated with the following parameters: *α*_*TD*_ = 0.01, *α*_*MB*_ = 0.50, *γ* = 0.93, *k* = 0.50. RPEs were extracted on test trials and visualised in fig. 4.

### 6.7 Mikhael et al. (2022) simulation

Finally, we compared dual-process RPEs with dopamine signals in Mikhael et al. (2022). In the Mikhael et al. (2022) experiment, subjects viewed a VR track akin to Kim et al. (2020) except that the sensory features were progressively darkened on a subset of trials. The experiment further incorporated speed-manipulations.

Environments were formalised with feature-based function approximation. Each state was initially encoded as a one-hot feature vector. To generate state uncertainty, feature vectors were passed through a Gaussian filter parameterised by 𝒩(*s*_*t*_, *σ*_*t*_). The mean of the Gaussian *s*_*t*_ was always the true-state at time *t*, while the standard deviation *sigma*_*t*_ was time dependent, and set differently in each experimental condition (see below). Lost probability mass (i.e. mass that was pushed beyond the boundaries of the environment due to Gaussian filtering) was reassigned to the nearest boundary state ensure simplex feature distributions.

In fig. 5C, we simulated a dual-process agent on the VR track experiment in Mikhael et al. (2022). The agent was simulated according to eqs. 5—7, but with value estimates constructed according to eq. (11) and eq. (12) to account for state uncertainty. The agent was tested on a linear track with *N* = 87 states with a goal state that delivered a reward *r* = 1.0 at one end of the track. In the bright condition, the standard-deviation in the Gaussian filter was constant at *σ*_*t*_ = 4. In the darkening condition, the standard-deviation was drawn from a rescaled exponential function with the minimum value *σ*_*min*_ = 4 and a maximum value *σ*_*max*_ = 24, which reproduced the assumptions about state uncertainty during sensory darkening in Mikhael et al. (2022). In the standard-speed condition, the agent moved with the stepsize *stepsize*_*std*_ = 1, whereas in the fast-speed condition, it moved with the stepsize *stepsize*_*fast*_ = 2. The agent was simulated with the following parameters: *α*_*TD*_ = 0.01, *α*_*MB*_ = 0.50, *γ* = 0.93, *k* = 0.50. It was first trained on the task in the bright, standard-speed condition for *T* = 150 trials. It then performed one test trial in each combination of brightness (bright-vs-dark) and speed (standard-vs-fast) conditions. The RPE in each state and each test condition was extracted and visualised in fig. 5C.

